# Characterization of a novel mouse model for Fuchs Endothelial Corneal Dystrophy

**DOI:** 10.1101/2023.10.06.561253

**Authors:** Subashree Murugan, Viviane Souza de Campos, Sachin Anil Ghag, Matthew Ng, Rajalekshmy Shyam

**Affiliations:** Vision Science Program, School of Optometry, Indiana University Bloomington, Indiana; Department of Biology, Indiana University Bloomington, Indiana

**Keywords:** FECD, corneal endothelial dystrophy, Col8a2, Slc4a11, guttae

## Abstract

**Purpose:** Fuchs Endothelial Corneal Dystrophy (FECD) is a progressive blinding disorder prevalent in 4% of Americans over 40. Corneal transplantation is the standard treatment. Animal models with partial FECD features exist, but a model encompassing all the major disease characteristics is desirable to improve the understanding of the pathogenesis and to identify signaling pathways involved in the disease onset and progression. Such an animal model can be helpful to develop intervention strategies. Here, we developed a mouse model that recapitulates all the features of FECD.

**Method:** Loss of function mutations in *Slc4a11* and a knock-in mutation in *Col8a2* (Q455K) are implicated in FECD. Mice with Slc4a11 and Col8a2 mutations in C57BL/6J background were crossed to generate double mutant mice at F2 generation. At five weeks of age, a subset of the animals were fed tamoxifen-enriched chow or standard chow for two weeks, followed by standard chow. Corneal thickness, endothelial cell density, and guttae were measured at 5 (baseline) and 16 weeks of age. Corneas collected from mice at 16 weeks were stained for tight and adherens junctions, and reactive oxygen species. A lactate assay was performed to evaluate the endothelial pump function.

**Results:** The double mutant tamoxifen-fed mice showed increased corneal thickness, decreased endothelial cell density, presence of guttae, and elevated stromal lactate levels. The endothelial cells showed altered morphology with disrupted adherens junctions and elevated ROS.

**Conclusion:** Overall, this mouse model recapitulates all the important phenotypic features associated with FECD.

## Introduction

The corneal endothelium is a non-regenerative ^1–3^, metabolically active monolayer that maintains corneal transparency through stromal deturgescence ^4–7^. Dysfunctions and degenerative changes in the endothelium result in bilateral non-inflammatory disorders known as corneal endothelial dystrophies^8,9^. Fuchs Endothelial Corneal Dystrophy (FECD) is a progressive blinding disorder prevalent in 4% of Americans over 40 years of age ^7,9,10^. The characteristic features of FECD include endothelial cell loss, guttae (extracellular matrix excrescences) formation, Descemet’s membrane thickening^11^, polymegethism (enlarged cell size), pleomorphism (change in cell morphology)^12,13^, and corneal edema, eventually leading to vision loss ^14^. Corneal transplantation is the most common treatment option available ^15–17^. Disadvantages of this treatment include graft rejection, a reduced survival rate of 5 years for the transplanted tissue, and secondary ocular complications like glaucoma and retinal detachment ^15^.

FECD progresses longitudinally with little to no visual deficits at the initial stage. Increased guttae formation is the first sign of disease^11^, followed by corneal endothelial cell loss and edema. FECD is a polygenic disease^7,15,18,19^ and has associations with several genes such as *TCF4* ^20,21^, *COL8A2* ^22–24^, *SLC4A11* ^25,26^, *ZEB1* ^27–29^ and *LOXHD1* ^30,31^. Most studies of FECD pathogenesis are conducted on end-stage human tissues and corneal endothelial cell lines ^32–35^. While informative, they provide a limited understanding of the disease progression.

Animal models can circumvent this problem; however, the prevalent mouse models for FECD represent only certain disease features. For instance: the *Col8a2* Q455K mouse, which contains an autosomal dominant mutation implicated in early-onset FECD, shows the presence of guttae and corneal endothelial cell loss but no corneal edema ^36^. Solute carrier family 4, member 11 (SLC4A11), an inner mitochondrial membrane protein, is highly expressed in the corneal endothelium, and the loss of function of this protein is associated with Congenital Hereditary Endothelial Dystrophy ^37,38^ and late-onset FECD ^26^. The transcript levels of this gene are reduced in the early-onset FECD mouse model ^25^. The *Slc4a11*^*-/-*^ mouse shows corneal edema and endothelial cell loss but no guttae^37^. While the *Slc4a11*^*-/-*^ and Col8a2 Q455K mouse models exhibit corneal endothelial dysfunctions, these mice are otherwise healthy with no developmental abnormalities ^36–38^.

Mitochondrial dysfunction associated with elevated oxidative stress is implicated in FECD pathogenesis ^39–44^ and in *Slc4a11*^*-/-*^ mice^45,46^. In a highly metabolic cell layer such as the corneal endothelium ^47^, which has the second highest density of mitochondria in the human body ^48^, the loss of *Slc4a11* can help identify the role of oxidative stress and mitochondrial dysfunctions on FECD onset and progression. Extracellular matrix (ECM)-associated changes evidenced by guttae, and Descemet’s membrane thickness are present in FECD ^11,16,33^ and Col8a2 Q455K mice ^24,36^. However, the role of the ECM in FECD disease pathogenesis is underexplored. As the animal models with individual mutations phenocopy only partial features of FECD, we hypothesized that the double mutant mice containing the loss of *Slc4a11* and *Col8a2* Q455K might provide a better understanding of the roles of oxidative stress and ECM-associated changes in FECD disease pathogenesis.

In this study, we characterized this double mutant mouse model and showed that it phenocopies all the major features of FECD, including guttae, corneal endothelial cell loss, and corneal edema. This mouse model also shows evidence of the interplay between elevated oxidative stress and extracellular matrix changes toward FECD progression.

## Materials and Methods

### Generation of transgenic mice and genotyping

All animal procedures were approved by the Institutional Animal Care and Use Committee (IACUC) at the Indiana University and adhered to the Association for Research in Vision and Ophthalmology (ARVO) statement for the use of animals in Ophthalmic and Vision research. A transgenic mouse containing two FECD mutations was generated by crossing the tamoxifen(Tm)-inducible knockdown of *Slc4a11* (*B6*.*Slc4a11*^*Flox/Flox*^/*Rosa*^*Cre-ERT2*/*Cre-ERT2*^) with Col8a2 (Q455K) knock-in (*B6*.Col8a2^*ki/ki*^) mice **(Figure 1A)**.The *Slc4a11*^*Flox/Flox*^/*Rosa*^*Cre-ERT2*/*Cre-ERT2*^ was a kind gift from Joseph Bonanno lab at the Indiana University School of Optometry^37^. Heterozygous mice with *Col8a2* knock-in mutation were purchased from the Jackson Laboratory (strain # 029749, Bar Harbor, ME). Both mice were in C57BL/6J (B6) background.

**Figure 1:**
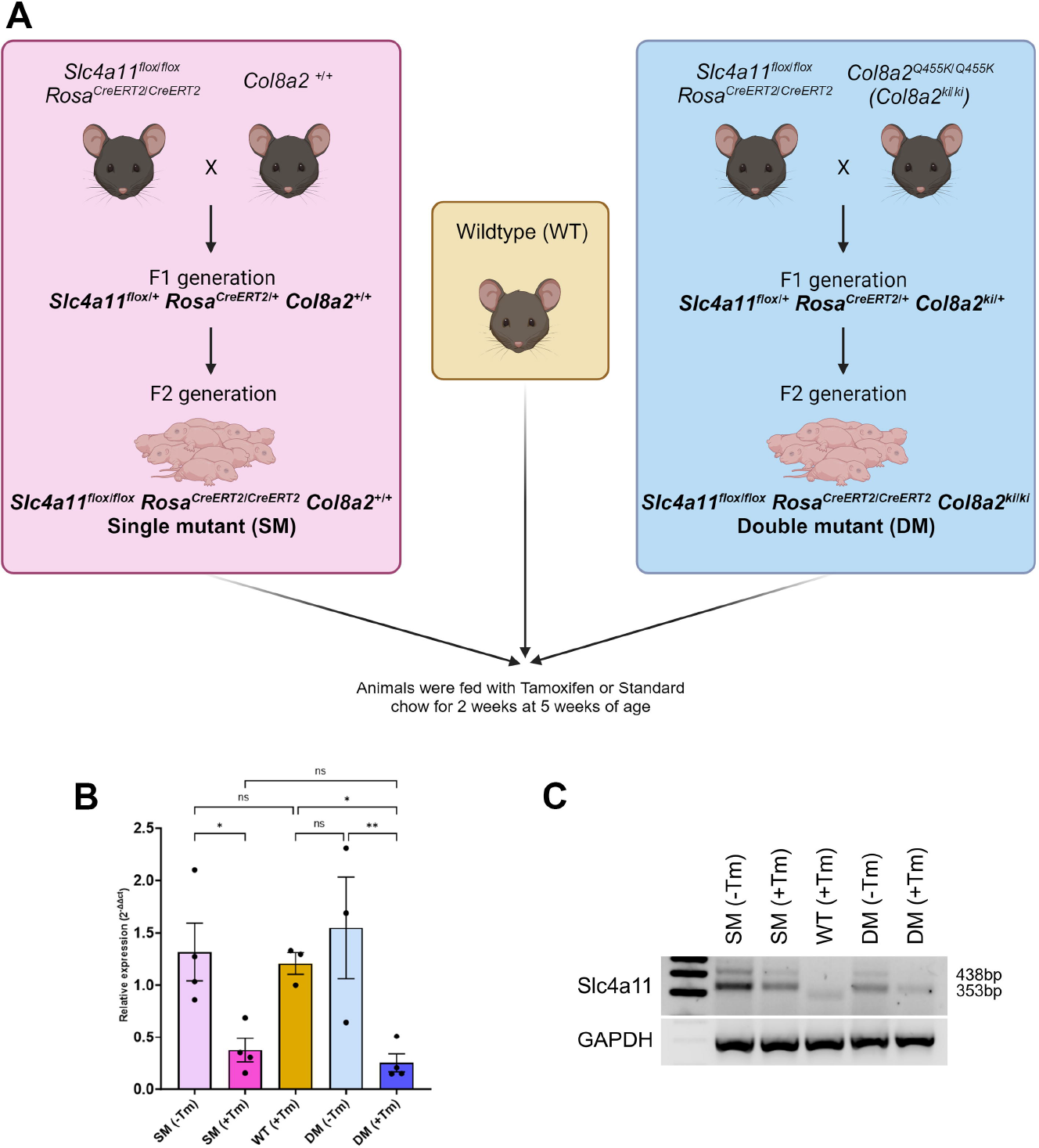
Generation of transgenic mice and verification of the *Slc4a11* knockdown. **A**. Schematic of the gene animals. The Tm-inducible knockdown of *Slc4a11* mouse (*B6*.*Slc4a11*^*Flox/Flox*^/*Rosa*^*Cre-ERT2*/*Cre-ERT2*^) was crossed either with Col8a2 wildtype (*B6*.*Col8a2*^*+/+*^) or Col8a2 (Q455K) knockin (*B6*.*Col8a2*^*ki/ki*^) animals to produce litters with the genotype *B6*.*Slc4a11*^*Flox/Flox*^/*Rosa*^*Cre-ERT2*/*Cre-ERT2*^/*Col8a2*^*+/+*^ or *B6*.*Slc4a11*^*Flox/Flox*^/*Rosa*^*Cre-ERT2*/*Cre-ERT2*^/*Col8a2*^*ki/ki*^ referred to as Single mutants (SM) and Double mutants (DM), respectively. The animals were either fed with tamoxifen (Tm) for 2 weeks at five weeks of age - SM(+Tm) or DM(+Tm), or fed with standard chow - SM(-Tm) or DM(-Tm). Age-matched wildtype animals were also fed with Tm for 2 weeks. **B**. Quantification of *Slc4a11* transcripts using kidney lysates from experimental SM(+Tm) or DM(+Tm), control SM(-Tm) or DM(-Tm), and wildtype (WT) animals at 16 weeks of age (n=3 to 4 replicates). **C**. cDNA was used to perform PCR to confirm the knockdown of Slc4a11 (floxed band-438bp; wildtype band-353bp) with GAPDH as a loading control. Mean + standard deviation (SD) ns-not significant, *P<0.05, ** P<0.001, ****P<0.0001. (1-way ANOVA with Uncorrected Fisher’s LSD multiple comparisons). *B6*.*Slc4a11*^*Flox/Flox*^/*Rosa*^*Cre-ERT2*/*Cre-ERT2*^/Col8a2^*+/+*^ fed with Tm – SM(+Tm), *B6*.*Slc4a11*^*Flox/Flox*^/*Rosa*^*Cre-ERT2*/*Cre-ERT2*^/Col8a2^*+/+*^ fed with normal chow – SM(-Tm), *B6*.*Slc4a11*^*Flox/Flox*^/*Rosa*^*Cre-ERT2*/*Cre-ERT2*^/Col8a2^ki/ki^ fed with Tm – DM(+Tm), *B6*.*Slc4a11*^*Flox/Flox*^/*Rosa*^*Cre-ERT2*/*Cre-ERT2*^/Col8a2^ki/ki^ fed with Tm – DM(-Tm)

The breeding scheme is outlined in Figure 1A. Briefly, we crossed *B6*.*Col8a2*^*+/+*^ mice with *B6*.*Slc4a11*^*Flox/Flox*^/*Rosa*^*Cre-ERT2*/*Cre-ERT2*^ mice to obtain *B6*.*Slc4a11*^*Flox/Flox*^/*Rosa*^*Cre-ERT2*/*Cre-ERT2*^/Col8a2^*+/+*^ at the F2 generation hereafter referred to as Single Mutants (SM). Similarly, we crossed *B6*.*Slc4a11*^*Flox/Flox*^/*Rosa*^*Cre-ERT2*/*Cre-ERT2*^ with *B6*.Col8a2^*ki/ki*^ animals to obtain *B6*.*Slc4a11*^*Flox/Flox*^/*Rosa*^*Cre-ERT2*/*Cre-ERT2*^/Col8a2^*ki/ki*^ at the F2 generation hereafter referred to as Double Mutants (DM). SM(+Tm) and DM(+Tm) were maintained on Tm chow (#TD.130859, Envigo, West Lafayette, IN) from five weeks of age until seven weeks of age. After this, these animals were maintained in standard chow until they were 16 weeks of age. Controls included SM(-Tm) and DM(-Tm) animals not fed Tm chow and wildtype animals fed with Tm chow. The mice were genotyped by sending ear punch samples for automated genotyping (Transnetyx, Cordova, TN).

### Total RNA isolation, cDNA synthesis and PCR

To test the knockdown of *Slc4a11* transcript following Tm, mouse kidneys from animals at 16 weeks of age were isolated from each group to isolate total RNA using the Trizol method. The kidney samples were homogenized in 1 mL of Trizol (Ambion, catalog #15596018) using a handheld homogenizer for 10 minutes. 200 µl of chloroform was added to the homogenate and mixed vigorously by hand for 15 seconds. The samples were centrifuged at 12,000g for 15 minutes at 4°C, after a brief incubation of 5 minutes at room temperature. The aqueous layer was collected in a new tube and mixed with an equal volume of isopropanol by inversion. The samples were centrifuged at 12,000g for 20 minutes at 4°C, after 10 minutes of incubation at room temperature. The supernatant was removed, and the pellet was washed with 1ml 75% ethanol and vortexed for 5 seconds. The samples were centrifuged at 13,000g for 10 minutes at 4°C. The pellets were air dried at room temperature and suspended in 50µl of DNase/RNase free water. The RNA was stored at −80°C until further clean-up using RNeasy Mini kit (Qiagen, catalog #74104) manufacturer’s protocol.

To synthesize cDNA from total RNA, a High-Capacity RNA-to-cDNA kit (Applied Biosystems, catalog #4387406) was used. Reverse transcription (RT) reaction mix was prepared by mixing 10µl of the 2X RT buffer mix, 1µl of RT enzyme mix, total RNA of 2µg and nuclease-free water up to 20µL per reaction. The RT reaction mix was aliquoted into PCR tubes and incubated at 37°C for 60 minutes. The reaction was stopped by heating to 95°C for 5 mins and held at 4°C.

The cDNA was used to perform real-time PCR using the following primers for *Slc4a11*: Forward - TCTGGACTTCAACGCCTTCT and Reverse - GCACAAACGTGATGGAAATG and Col8a2: Forward - ATTCGAGGAGACCAAGGGCCTAAT and Reverse - AAGTGAGCACTGCAGTAAAGGCTG. Real-time PCR was performed using the commercially available SsoAdvanced Universal SYBR Green Supermix following manufacturer’s instructions (BIO-RAD, catalog #1725271).The cDNA was amplified using Platinum Direct PCR Universal Master mix (Invitrogen, catalog #A44647100) manufacturer’s protocol to perform regular PCR, followed by gel electrophoresis.

### Optical Coherence Tomography and Heidelberg Retinal Tomography 3 – Rostock Cornea Module

Anterior segment - Optical Coherence Tomography (AS-OCT) (iVue100 Optovue, Inc., Fremont, CA, USA) was performed for pachymetry measurements and Heidelberg Retinal Tomography 3 – Rostock Cornea Module (HRT3-RCM) (Heidelberg Engineering Inc., Franklin, MA, USA) for *in vivo* assessment of the corneal endothelium was performed. Corneal thickness measurements and endothelial assessment were done before Tm treatment at five weeks of age (baseline) and at 16 weeks of age. The mice were anesthetized using a solution containing 100mg/kg of ketamine and 10mg/kg of xylazine, administered intraperitoneally. The ocular surface was kept moist using a saline solution during AS-OCT. Horizontal scans of the central cornea were taken using the widest pupil diameter as a reference. HRT3-RCM was done at a 400 x 400 µm field of view to obtain enface images of the corneal endothelium. Genteal eye gel (Alcon) was used to lubricate the corneal surface and as a coupling agent between the objective lens and the mouse cornea. Section scans at the corneal endothelial plane were acquired and used for assessing the number of cells and guttae.

### Immunofluorescence

Corneal cups were dissected from fresh mouse eyes, washed twice with 1X Phosphate Buffer solution (PBS), and placed in a 96-well plate. The tissue was fixed for 10 mins with 4% paraformaldehyde in 1x PBS at room temperature, washed with 1X PBS twice for 5 mins, and permeabilized and blocked with 0.5% Triton X-100 (Fisher Scientific, catalog # BP151-500) and 5% Normal donkey serum (Jackson Immuno Research, catalog #017-000-121) in 1X PBS for 30 minutes at room temperature. The tissues were incubated at 4°C overnight with the following primary antibodies diluted in the antibody diluent containing 0.1% Triton X-100 and 2% Normal donkey serum: Mouse anti-ZO-1(Fisher Scientific, catalog #339100) (1:100), Rabbit anti-N-Cadherin (Cell Signaling Technology, catalog #13116s) (1:100) and Rabbit anti-Nitrotyrosine (Life Technologies Corporation, catalog #A21285) (1:100). The tissue was washed with 1X PBS and incubated with Goat anti-Mouse IgG, Alexa Fluor™ 594 (Thermo Fisher, catalog #A11032) and/or Goat anti-Rabbit IgG; Alexa Fluor™ 488 (Thermo Fisher, catalog #A11034) secondary antibodies at 1:100 dilution for an hour at room temperature. The corneal cups were washed with 1X PBS and flat-mounted using a drop of Prolong Glass Antifade Mountant with NucBlue (Invitrogen, catalog #P36985). The tissues were imaged using a Zeiss Apotome2 microscope (Carl Zeiss, White Plains, NY, USA). One cornea from at least three animals were used for each staining condition. For Nitrotyrosine stained corneas, three images of central cornea were taken at similar laser intensity, gain and exposure time. The images were analyzed using ImageJ to measure the mean fluorescence intensity in each image. The average staining intensity was used for statistical analysis.

### Lactate assay

Whole corneas were isolated from mouse eyes, weighed before pulverizing using liquid nitrogen and homogenized in 30 µl of PBS using a motor and pestle. The supernatant was separated from the pellet by centrifugation at 15,000 G for 15 mins at 4C. The supernatant was used to measure the lactate levels using a lactate assay kit-WST (Dojindo Molecular Technologies, catalog #L256) per the manufacturer’s instructions.

### Statistical analysis

The quantification data was analyzed using GraphPad Prism Software (v.10.0.2, Boston, MA, USA). The bar graphs with error bars are representative of the mean and standard deviation. One-way ANOVA with Uncorrected Fisher’s LSD multiple comparisons was performed to compare the different groups and determine statistical significance (p-value of > 0.05 was considered significant).

### Illustrations

All illustrations were created using BioRender.com.

## Results

### Confirmation of *Slc4a11* knockdown and *Col8a2* knock-in in the double mutant animals

To assess the efficacy of Tm-mediated knockdown of *Slc4a11* expression, we evaluated *Slc4a11* transcript levels in SM(+Tm) and DM(+Tm) compared to SM (-Tm) and DM (-Tm). In addition to the corneal endothelium, *Slc4a11* is highly expressed in the kidneys^49,50^. Therefore, we used kidney lysate to confirm Slc4a11 knockdown. The relative expression of *Slc4a11* was reduced in the SM(+Tm) and DM(+Tm) mutant animals when compared to SM(-Tm), DM(-Tm), and wildtype animals **(Figure 1B)**. Gel electrophoresis of PCR products showed little to no expression of the floxed allele in the SM(+Tm) and DM(+Tm) **(Figure 1C)**. At the same time, the animals not fed with Tm, SM(-Tm) and DM(-Tm) showed the presence of the floxed and the wildtype allele. *Slc4a11* WT allele showed bands at 353 bp and floxed allele at 438 bp. The wildtype animals displayed only the wildtype band at 353 bp **(Figure 1C)**.

### Elevated corneal edema and lactate content in the double mutant animals

One of the characteristic features of FECD is the increase in corneal thickness due to stromal edema. We conducted pachymetry measurements using OCT on the animals at five weeks (baseline) and 16 weeks of age to determine any changes in corneal thickness. We observed a significant difference in corneal thickness in the SM(+Tm) and DM(+Tm) animals at 16 weeks of age when compared to their baseline measurements **(Figure 2A-B)**. On average, the SM(+Tm) and the DM(+Tm) showed approximately 50 µm increase in corneal thickness. No significant elevation in corneal thickness was observed in the wildtype animals, SM(-Tm) and DM(-Tm) **(Figure 2A-B)**.

**Figure 2:**
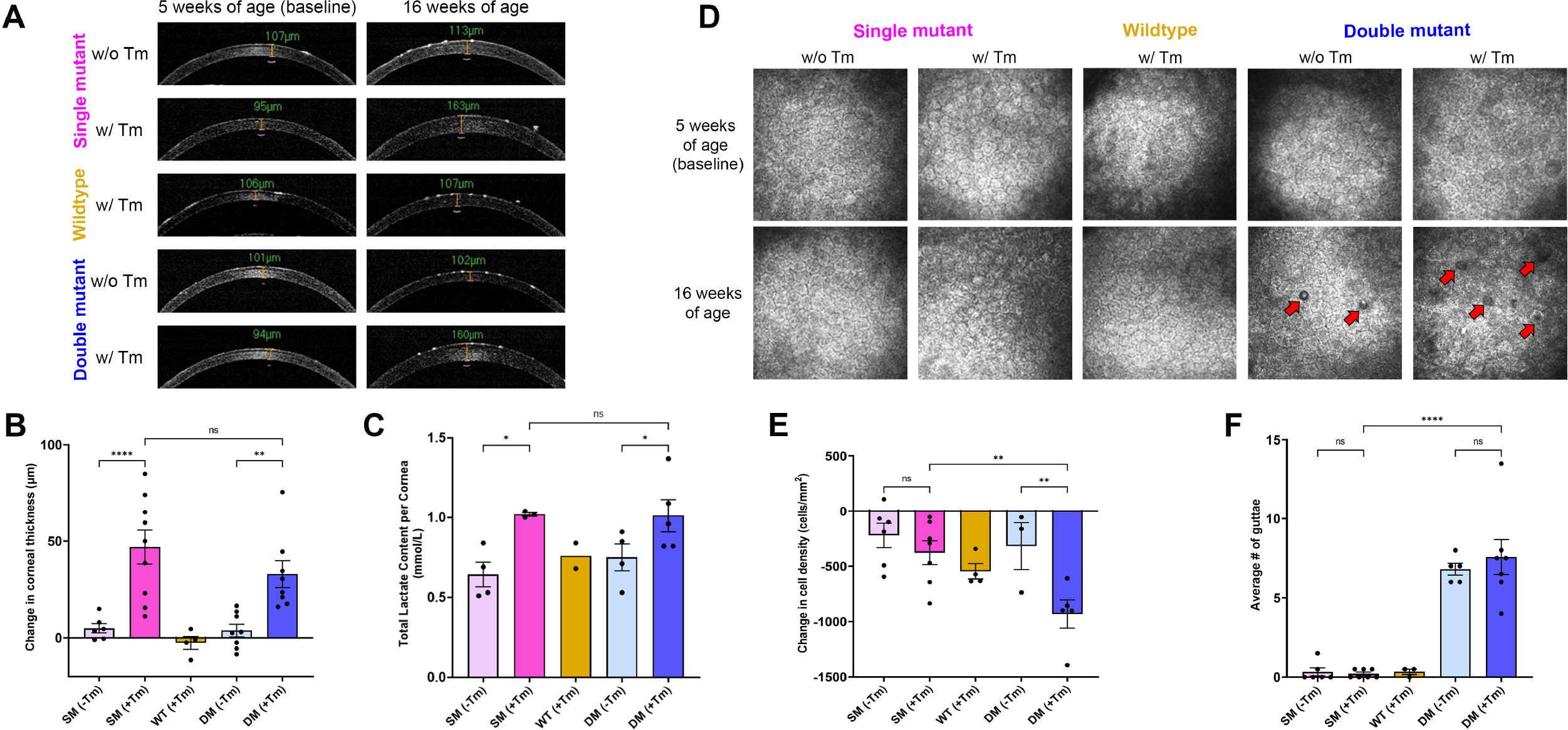
Phenotypic assessment. **A**. Representative OCT images of SM(-Tm), SM(+Tm), WT, DM(-Tm), and DM(+Tm) animals at 5 weeks of age (baseline) and at 16 weeks of age. **B**. Quantification of the change in corneal thickness at 16 weeks of age from baseline (n=4 to 9 eyes) **C**. Quantification of lactate content (n=2 to 5 eyes). **D**. Representative HRT3-RCM images of SM(-Tm), SM(+Tm), WT, DM(-Tm), and DM(+Tm) corneal endothelium (FOV 400um) at 5 weeks of age (baseline) and 16 weeks of age; red arrows point to guttae. **E**. Quantification of the change in the endothelial cell density at 16 weeks of age from baseline (n=3 to 7 eyes). **F**. Quantification of the average number of guttae in the right and left eyes (n=3 to 7 eyes); Mean + standard deviation (SD) ns-not significant, *P<0.05, ** P<0.001, ****P<0.0001. (1-way ANOVA with Uncorrected Fisher’s LSD multiple comparisons). *B6*.*Slc4a11*^*Flox/Flox*^/*Rosa*^*Cre-ERT2*/*Cre-ERT2*^/Col8a2^*+/+*^ fed with Tm – SM(+Tm), *B6*.*Slc4a11*^*Flox/Flox*^/*Rosa*^*Cre-ERT2*/*Cre-ERT2*^/Col8a2^*+/+*^ fed with normal chow – SM(-Tm), *B6*.*Slc4a11*^*Flox/Flox*^/*Rosa*^*Cre-ERT2*/*Cre-ERT2*^/Col8a2^ki/ki^ fed with Tm – DM(+Tm), *B6*.*Slc4a11*^*Flox/Flox*^/*Rosa*^*Cre-ERT2*/*Cre-ERT2*^/Col8a2^ki/ki^ fed with Tm – DM(-Tm)

Corneal edema is directly proportional to the lactate accumulation resulting from the decrease in the efflux of lactate from the stroma ^51^. Consistent with the pachymetry results, we observed an elevation in the lactate levels in the SM(+Tm) and double DM(+Tm) when compared to the controls. In both SM(+Tm) and DM(+Tm) animals, we observed nearly 50% elevation of the lactate levels when compared to the controls **(Figure 2C)**.

### Decreased corneal endothelial cell count and increased guttae formation in the double mutant animals

FECD is characterized by the loss of corneal endothelial cells and an elevation in guttae formation^5,7,13,52^. We assessed the corneal endothelium using HRT3-RCM **(Figure 2D-F)** to determine changes in corneal endothelial density and guttae. Compared to the baseline measurements at five weeks of age, we saw a significant decrease in cell density with less than 1000 cells/mm^2^ in the DM (+Tm) animals. On the other hand, control animals lost approximately 500 cells/mm^2^, consistent with the age-related loss of corneal endothelial density^53–56^.

Interestingly, the guttae numbers were comparable between DM(+Tm) and DM(-Tm), with both groups showing approximately ten guttae per captured image area. As expected, SM(+Tm) and wildtype animals showed very few guttae in our analysis.

FECD is characterized by the loss of corneal endothelial cells and an elevation in guttae formation ^5,7,13,52^. We assessed the corneal endothelium using HRT3-RCM **(Figure 2D-F)** to determine changes in corneal endothelial density and guttae. Compared to the baseline measurements at five weeks of age, we saw a significant decrease in cell density with less than 1000 cells/mm^2^ in the DM(+Tm) animals **(Figure 2D-E)**. On the other hand, control animals lost approximately 500 cells/mm^2^ **(Figure 2D-E)**, consistent with the age-related loss of corneal endothelial density seen in humans ^3,56–58^. Thus, we were able to exclude the effect of Tm on endothelial cell loss. Interestingly, the guttae numbers were comparable between DM(+Tm) and DM(-Tm), with both groups showing approximately ten guttae per captured image area **(Figure 2D**,**F)**. As expected, SM(+Tm) and wildtype animals showed very few guttae in our analysis **(Figure 2D,F)**.

### Changes in the adherens junction and hexagonality of the endothelial cells in the double mutants

N-Cadherin is a major component of the adherens junctions in the corneal endothelial cells^59,60^. Cell adhesion is essential in the maintenance of corneal endothelial cell integrity and function^61^. We, therefore, checked if there were disruptions in the N-Cadherin expression patterns in the animals. Immunofluorescence of the corneal cups stained with an antibody against N-Cadherin revealed highly disrupted adherens junction **(Figure 3A)** in the SM(+Tm) and DM(+Tm). In contrast, we observed well-organized N-Cadherin staining patterns in the SM(-Tm), DM(-Tm), and WT animals.

**Figure 3:**
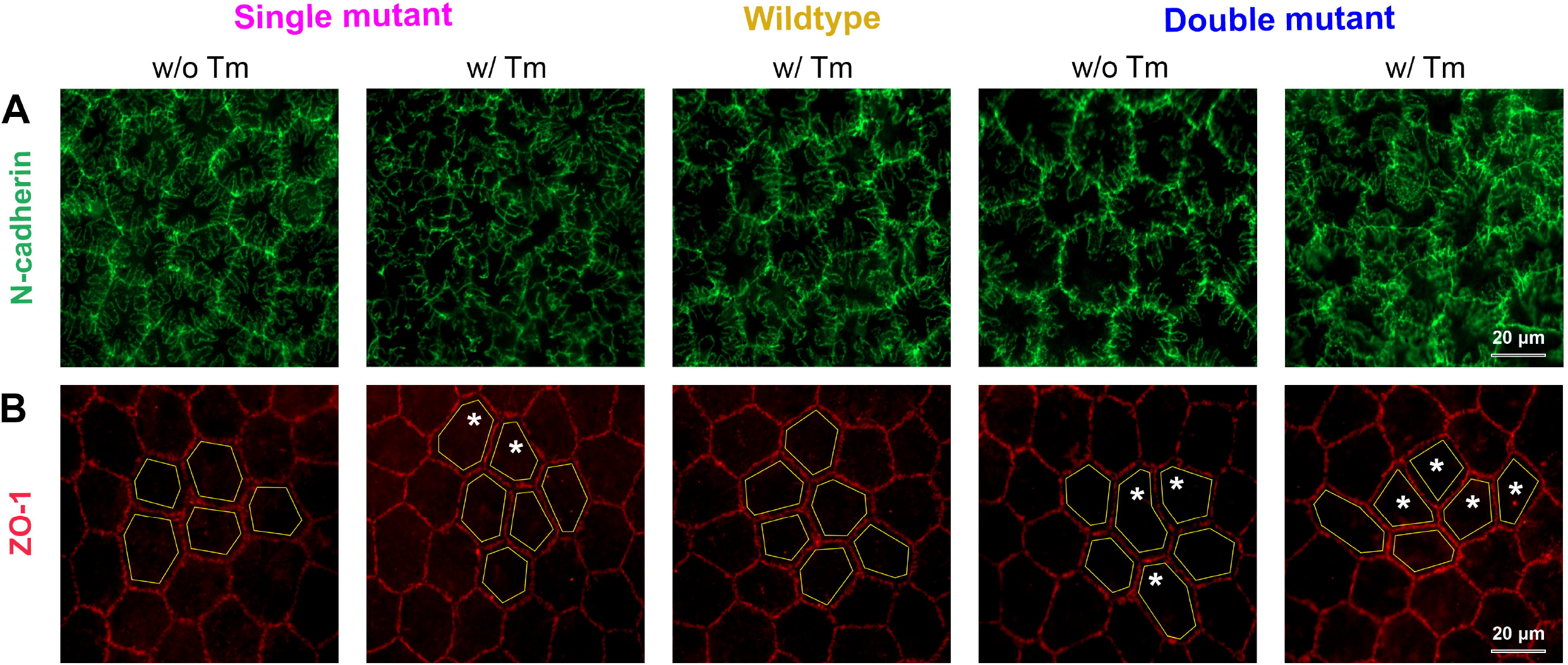
Evaluation of Adherens junctions and cell morphology in corneal endothelial flatmounts. **A**. Representative images of N-cadherin staining to visualize adherens junction in the corneal endothelium of SM(-Tm), SM(+Tm), WT, DM(-Tm), and DM(+Tm) animals at 16 weeks of age (n=3 eyes). **B**. Representative images of ZO-1 staining to visualize the endothelial cell morphology and hexagonality in SM(-Tm), SM(+Tm), WT, DM(-Tm), and DM(+Tm) animals at 16 weeks of age (n=3 eyes). Yellow lines are presented to visualize the hexagonality of the endothelial cells and white asteriks indicate the pleomorphic cells. Scale bar 20 mm. *B6*.*Slc4a11*^*Flox/Flox*^/*Rosa*^*Cre-ERT2*/*Cre-ERT2*^/Col8a2^*+/+*^ fed with Tm – SM(+Tm), *B6*.*Slc4a11*^*Flox/Flox*^/*Rosa*^*Cre-ERT2*/*Cre-ERT2*^/Col8a2^*+/+*^ fed with normal chow – SM(-Tm), *B6*.*Slc4a11*^*Flox/Flox*^/*Rosa*^*Cre-ERT2*/*Cre-ERT2*^/Col8a2^ki/ki^ fed with Tm – DM(+Tm), *B6*.*Slc4a11*^*Flox/Flox*^/*Rosa*^*Cre-ERT2*/*Cre-ERT2*^/Col8a2^ki/ki^ fed with Tm – DM(-Tm)

Loss of hexagonality and cell size changes are associated with FECD. We carried out ZO-1 staining to assess changes to cell morphology. A regular pattern of cellular arrangements and cell size was evident in the corneal endothelium of wildtype and SM(-Tm) animals based on ZO-1 staining **(Figure 3B)**. However, in both DM(+Tm) and DM(-Tm) animals, we observed comparable changes in cell morphology, including the loss of hexagonal shape and cell size changes **(Figure 3B)**.

### Elevated oxidative stress in the double mutant corneal endothelium

Nitrotyrosine is a marker for elevated oxidative stress^62,63^. Since FECD pathogenesis is associated with increased oxidative stress, we evaluated if there were changes in nitrotyrosine levels in the corneal endothelium of the double mutants. As expected, the SM(-Tm) and the wildtype animals showed minimal nitrotyrosine staining. Increased staining was evident in SM (+Tm), confirming previous findings^37^. Interestingly, we observed a nearly 4-fold increase in nitrotyrosine staining in the DM (+Tm) compared to the SM (+Tm) **(Figures 4A and B)**.

**Figure 4:**
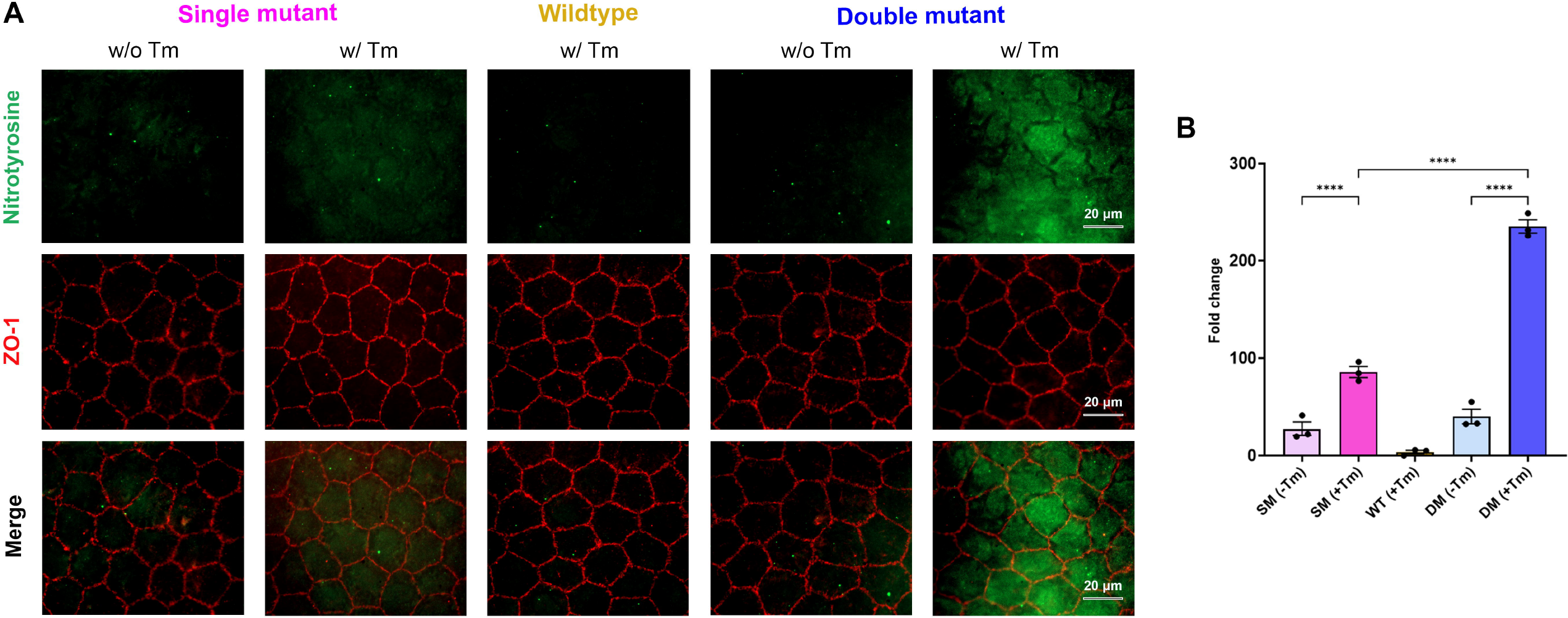
Evaluation of oxidative stress in corneal endothelium flatmounts. **A**. Representative nitrotyrosine, ZO-1 and merge images to visualize the reactive oxygen species in the corneal endothelium SM(-Tm), SM(+Tm), WT, DM(-Tm), and DM(+Tm) animals at 16 weeks of age. **B**. Quantification of the fold change in mean fluorescence from A (n=3 eyes). Scale bar 20um. Mean + standard deviation (SD) ****P<0.0001. (1-way ANOVA with Uncorrected Fisher’s LSD multiple comparisons). *B6*.*Slc4a11*^*Flox/Flox*^/*Rosa*^*Cre-ERT2*/*Cre-ERT2*^/Col8a2^*+/+*^ fed with Tm – SM(+Tm), *B6*.*Slc4a11*^*Flox/Flox*^/*Rosa*^*Cre-ERT2*/*Cre-ERT2*^/Col8a2^*+/+*^ fed with normal chow – SM(-Tm), *B6*.*Slc4a11*^*Flox/Flox*^/*Rosa*^*Cre-ERT2*/*Cre-ERT2*^/Col8a2^ki/ki^ fed with Tm – DM(+Tm), *B6*.*Slc4a11*^*Flox/Flox*^/*Rosa*^*Cre-ERT2*/*Cre-ERT2*^/Col8a2^ki/ki^ fed with Tm – DM(-Tm)

## Discussion

In this study, we characterized the double mutant mouse model encompassing all the disease phenotypes associated with FECD. One of the first signs of FECD is the presence of guttae, followed by corneal endothelial cell loss and corneal edema^11,16,33^. In the double mutant mouse, we observed guttae formation, corneal endothelial cell loss, and edema, all of which phenocopies the FECD in human patients **(Figure 5A)**. Pioneering studies using immortalized corneal endothelial cells and end-stage disease tissues showed that elevated oxidative stress is a cause of FECD progression^25,32–34^. However, the lack of *in-vivo* evidence limits the understanding of how oxidative stress can lead to the disease. The loss of *SLC4A11* elevates oxidative stress in the corneal endothelium^38,40,42–44^, and *Slc4a11*^*-/-*^ mice show elevated corneal edema associated with reactive oxygen species^64^. The effects of *Col8a2* Q455K KI mutation on oxidative stress are not well-studied. In the DM(+Tm), we observed corneal edema comparable to the SM(+Tm) levels, indicating that elevated oxidative stress can increase corneal thickness. Even though the DM(+Tm) showed nearly a 4-fold increase in oxidative stress staining, we did not observe a comparable elevation in corneal thickness in these animals. This finding suggests that further evaluation of corneal edema-oxidative stress dynamics is required.

**Figure 5:**
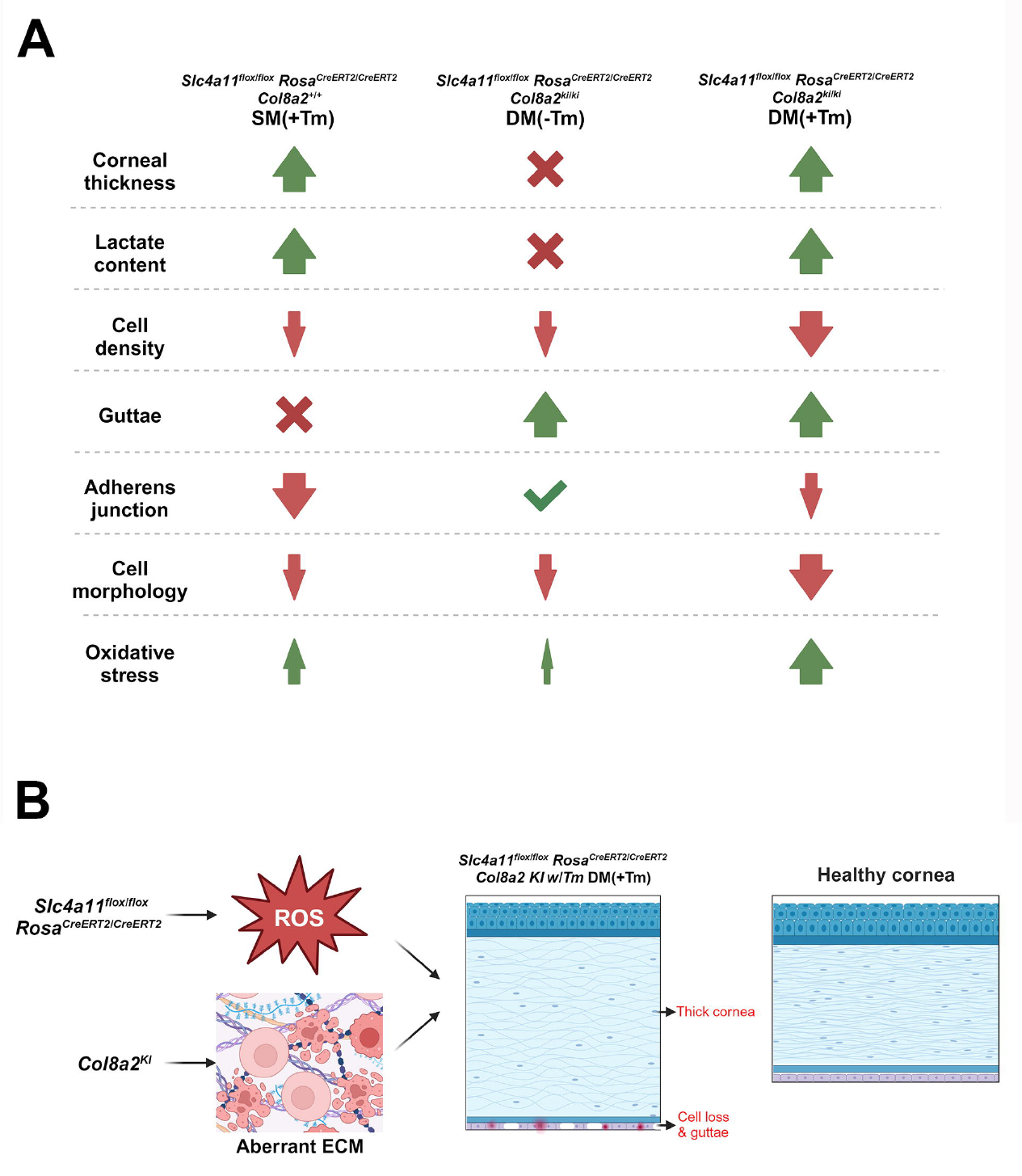
**A**. Summary of the characteristic features noted in the experimental and control animals in Fuchs Endothelial Corneal Dystrophy. The double mutant showed a significant reduction in cell density, N-cadherin expression, and cell morphology when compared to Slc4a11 knockdown and Col8a2 knock-in mutation. Guttae formation was comparable between Col8a2 Q455K knock-in mutants and the double mutant. Significant elevation in oxidative stress was evident in the double mutant when compared to Slc4a11 knockdown. **B**. Schematic showing the effect of the Slc4a11 knock down and Col8a2 knock in mutations in the *B6*.*Slc4a11*^*Flox/Flox*^/*Rosa*^*Cre-ERT2*/*Cre-ERT2*^ DM(+Tm) mouse model.

Central corneal guttae are one of the first features associated with FECD in humans. Progressive guttae are present in the Col8a2 Q455K mice ^36^, but not in the *Slc4a11*^*-/-*^ animals. While the DM(+Tm) showed guttae before corneal edema or corneal endothelial cell loss (data not shown), we did not observe a significant increase in the guttae numbers between the double mutant animals DM(+Tm) and the Col8a2 Q455K littermate DM(-Tm) controls. This finding indicates that elevated oxidative stress might not be causal in guttae formation. Future analysis of this mouse model will help identify the signaling pathways associated with guttae formation.

YAP/TAZ signaling pathways regulate cell shape changes ^65^. Polymegathism is a characteristic feature of FECD^12,13^. To identify whether the loss of YAP/TAZ pathways would result in corneal endothelial dysfunctions, Thomasy and colleagues used a knockout mouse model of *Wwrt1*, a primary YAP/TAZ pathway transducer^66^. This mouse model presented decreased corneal endothelial cell numbers, shape changes, and a softer Descemet’s Membrane, among the characteristic features of FECD. This study reveals the role of YAP/TAZ signaling in corneal endothelial homeostasis^66^. However, corneal thinning and the absence of guttae in the *Wwrt1*^*-/-*^ animal model indicate that the YAP/TAZ pathways may not alone be responsible for the FECD disease phenotypes. Whether the cell shape changes observed in the double mutant animals arise due to aberrant YAP/TAZ signaling can be evaluated in future studies.

Jurkunas group showed that the Ultraviolet A-associated DNA damage in FECD is associated with estrogen levels^67^. Their study was the first to reveal a connection between estrogen levels and FECD. Recently, a study from the Patel lab^68^ also found that estrogen-associated dysfunctions in human corneal endothelial cells are specific to the female sex. While the initial analysis of the double mutant animals did not reveal any sex-specific differences, we plan to determine the effects of estrogen-associated signaling pathways in FECD disease progression.

Corneal transplantation is the treatment option available for patients with FECD in the United States. Descemet stripping automated endothelial keratoplasty (DSAEK) is a prevalent form of posterior lamellar corneal transplantation^69^. However, this is an expensive procedure, costing over $12,000 per transplant. Graft-related complications arise approximately 5% of the time, and graft rejection is common in around 4% of the cases ^70^. Cell injection therapy is gaining traction as a corneal endothelial dystrophy treatment in Japan, with improved visual acuity after five years of the procedure ^71^. However, in their 5-year follow-up study, the authors indicate that guttae persist in the corneal endothelial junctions^72^. Since the effects of guttae on cell functions are yet to be understood, the long-term efficacy of this approach will require further evaluation. In this regard, the double mutant animal model can provide insights into understanding the changes to cellular dynamics associated with guttae formation.

## Conclusion

FECD is a prevalent corneal dystrophy with a global prevalence estimate of 7.33%^73^. The pathophysiology of this dystrophy is not entirely understood. The double Slc4a11 and Col8a2 mutant mouse model described here encompasses the two gene mutations reported in human patients. Characterization of the double mutant mouse model showed all the main features of FECD **(Figure 5B)**. FECD is a multifactorial disease with genetic and environmental causes. A limitation of the double mutant mouse model is that it does not represent all the causal factors associated with FECD. However, our analysis highlights that the double mutant is a significant advancement over the prevalent animal models **(Figure 5B)**. Future studies will evaluate this animal model to study the pathways involved in FECD disease onset and progression.

## Acknowledgements

We thank Dr. Joseph Bonanno, Dr. Mallika Valapala, and Dr. S.P. Srinivas (Indiana University Bloomington) for their valuable input throughout this project. We are grateful for the critical feedback on this manuscript from Dr. Krishnakumar Kizhatil (The Jackson Laboratory) and Dr. Elizabeth Zuniga-Sanchez (Baylor College of Medicine). This project was completed with the funding support from NIH R00 EY032974.

## Notes

**Grant support:** R00 EY032974, National Institutes of Health (NIH), 9000 Rockville Pike, Bethesda, Maryland 20892

### Competing Interest Statement

The authors have declared no competing interest.

## References

1. Feizi S. Corneal endothelial cell dysfunction: etiologies and management. Ophthalmol Eye Dis. 2018;10:2515841418815802. doi:10.1177/2515841418815802

2. Engelmann K, Böhnke M, Friedl P. Isolation and long-term cultivation of human corneal endothelial cells. Investigative Ophthalmology & Visual Science. 1988;29(11):1656–1662.

3. Joyce NC, Meklir B, Joyce SJ, Zieske JD. Cell cycle protein expression and proliferative status in human corneal cells. Investigative Ophthalmology & Visual Science. 1996;37(4):645–655.

4. Riley MV, Winkler BS, Starnes CA, Peters MI, Dang L. Regulation of corneal endothelial barrier function by adenosine, cyclic AMP, and protein kinases. Investigative Ophthalmology & Visual Science. 1998;39(11):2076–2084.

5. Bourne WM. Biology of the corneal endothelium in health and disease. Eye (Lond). 2003;17(8):912–918. doi:10.1038/sj.eye.6700559

6. Fischbarg J, Maurice DM. An update on corneal hydration control. Exp Eye Res. 2004;78(3):537–541. doi:10.1016/j.exer.2003.09.010

7. Wilson SE, Bourne WM. Fuchs’ dystrophy. Cornea. 1988;7(1):2–18.

8. Fuchs E. Dystrophia epithelialis corneae. Graefes Arhiv für Ophthalmologie. 1910;76(3):478–508. doi:10.1007/BF01986362

9. Schmedt T, Silva MM, Ziaei A, Jurkunas U. Molecular Bases of Corneal Endothelial Dystrophies. Exp Eye Res. 2012;95(1):24–34. doi:10.1016/j.exer.2011.08.002

10. Krachmer JH, Purcell JJ, Young CW, Bucher KD. Corneal endothelial dystrophy. A study of 64 families. Arch Ophthalmol. 1978;96(11):2036–2039. doi:10.1001/archopht.1978.03910060424004

11. Hogan MJ, Wood I, Fine M. Fuchs’ endothelial dystrophy of the cornea. 29th Sanford Gifford Memorial lecture. Am J Ophthalmol. 1974;78(3):363–383. doi:10.1016/0002-9394(74)90224-4

12. Polack FM. The posterior corneal surface in Fuchs’ dystrophy. Scanning electron microscope study. Invest Ophthalmol. 1974;13(12):913–922.

13. Waring GO, Rodrigues MM, Laibson PR. Corneal dystrophies. II. Endothelial dystrophies. Surv Ophthalmol. 1978;23(3):147–168. doi:10.1016/0039-6257(78)90151-0

14. Kannabiran C, Chaurasia S, Ramappa M, Mootha VV. Update on the genetics of corneal endothelial dystrophies. Indian J Ophthalmol. 2022;70(7):2239–2248. doi:10.4103/ijo.IJO_992_22

15. Matthaei M, Hribek A, Clahsen T, Bachmann B, Cursiefen C, Jun AS. Fuchs Endothelial Corneal Dystrophy: Clinical, Genetic, Pathophysiologic, and Therapeutic Aspects. Annu Rev Vis Sci. 2019;5:151–175. doi:10.1146/annurev-vision-091718-014852

16. Nanda GG, Alone DP. REVIEW: Current understanding of the pathogenesis of Fuchs’ endothelial corneal dystrophy. Mol Vis. 2019;25:295–310.

17. Zhang J, McGhee CNJ, Patel DV. The Molecular Basis of Fuchs’ Endothelial Corneal Dystrophy. Mol Diagn Ther. 2019;23(1):97–112. doi:10.1007/s40291-018-0379-z

18. Rosenblum P, Stark WJ, Maumenee IH, Hirst LW, Maumenee AE. Hereditary Fuchs’ Dystrophy. Am J Ophthalmol. 1980;90(4):455–462. doi:10.1016/s0002-9394(14)75011-1

19. Magovern M, Beauchamp GR, McTigue JW, Fine BS, Baumiller RC. Inheritance of Fuchs’ combined dystrophy. Ophthalmology. 1979;86(10):1897–1923. doi:10.1016/s0161-6420(79)35340-4

20. Mootha VV, Gong X, Ku HC, Xing C. Association and Familial Segregation of CTG18.1 Trinucleotide Repeat Expansion of TCF4 Gene in Fuchs’ Endothelial Corneal Dystrophy. Investigative Ophthalmology & Visual Science. 2014;55(1):33–42. doi:10.1167/iovs.13-12611

21. Wieben ED, Aleff RA, Tosakulwong N, et al. A Common Trinucleotide Repeat Expansion within the Transcription Factor 4 (TCF4, E2-2) Gene Predicts Fuchs Corneal Dystrophy. PLOS ONE. 2012;7(11):e49083. doi:10.1371/journal.pone.0049083

22. Biswas S, Munier FL, Yardley J, et al. Missense mutations in COL8A2, the gene encoding the α2 chain of type VIII collagen, cause two forms of corneal endothelial dystrophy. Human Molecular Genetics. 2001;10(21):2415–2423. doi:10.1093/hmg/10.21.2415

23. Gottsch JD, Sundin OH, Liu SH, et al. Inheritance of a novel COL8A2 mutation defines a distinct early-onset subtype of fuchs corneal dystrophy. Invest Ophthalmol Vis Sci. 2005;46(6):1934–1939. doi:10.1167/iovs.04-0937

24. Meng H, Matthaei M, Ramanan N, et al. L450W and Q455K Col8a2 knock-in mouse models of Fuchs endothelial corneal dystrophy show distinct phenotypes and evidence for altered autophagy. Invest Ophthalmol Vis Sci. 2013;54(3):1887–1897. doi:10.1167/iovs.12-11021

25. Vithana EN, Morgan PE, Ramprasad V, et al. SLC4A11 mutations in Fuchs endothelial corneal dystrophy. Human Molecular Genetics. 2008;17(5):656–666. doi:10.1093/hmg/ddm337

26. Riazuddin SA, Vithana EN, Seet LF, et al. Missense mutations in the sodium borate cotransporter SLC4A11 cause late-onset Fuchs corneal dystrophy. Hum Mutat. 2010;31(11):1261–1268. doi:10.1002/humu.21356

27. Gupta R, Kumawat BL, Paliwal P, et al. Association of ZEB1 and TCF4 rs613872 changes with late onset Fuchs endothelial corneal dystrophy in patients from northern India. Mol Vis. 2015;21:1252–1260.

28. Mehta JS, Vithana EN, Tan DTH, et al. Analysis of the Posterior Polymorphous Corneal Dystrophy 3 Gene, TCF8, in Late-Onset Fuchs Endothelial Corneal Dystrophy. Investigative Ophthalmology & Visual Science. 2008;49(1):184–188. doi:10.1167/iovs.07-0847

29. Riazuddin SA, Zaghloul NA, Al-Saif A, et al. Missense Mutations in TCF8 Cause Late-Onset Fuchs Corneal Dystrophy and Interact with FCD4 on Chromosome 9p. The American Journal of Human Genetics. 2010;86(1):45–53. doi:10.1016/j.ajhg.2009.12.001

30. Afshari NA, Igo RP, Morris NJ, et al. Genome-wide association study identifies three novel loci in Fuchs endothelial corneal dystrophy. Nature Communications. 2017;8. doi:10.1038/ncomms14898

31. Riazuddin SA, Parker DS, McGlumphy EJ, et al. Mutations in LOXHD1, a Recessive-Deafness Locus, Cause Dominant Late-Onset Fuchs Corneal Dystrophy. The American Journal of Human Genetics. 2012;90(3):533–539. doi:10.1016/j.ajhg.2012.01.013

32. Miyai T, Vasanth S, Melangath G, et al. Activation of PINK1-Parkin-Mediated Mitophagy Degrades Mitochondrial Quality Control Proteins in Fuchs Endothelial Corneal Dystrophy. Am J Pathol. 2019;189(10):2061–2076. doi:10.1016/j.ajpath.2019.06.012

33. Okumura N, Hashimoto K, Kitahara M, et al. Activation of TGF-beta signaling induces cell death via the unfolded protein response in Fuchs endothelial corneal dystrophy. Sci Rep. 2017;7(1):6801. doi:10.1038/s41598-017-06924-3

34. Yagi-Yaguchi Y, Yamaguchi T, Higa K, et al. Association between corneal endothelial cell densities and elevated cytokine levels in the aqueous humor. Sci Rep. 2017;7(1):13603. doi:10.1038/s41598-017-14131-3

35. Pan P, Weisenberger DJ, Zheng S, et al. Aberrant DNA methylation of miRNAs in Fuchs endothelial corneal dystrophy. Sci Rep. 2019;9:16385. doi:10.1038/s41598-019-52727-z

36. Jun AS, Meng H, Ramanan N, et al. An alpha 2 collagen VIII transgenic knock-in mouse model of Fuchs endothelial corneal dystrophy shows early endothelial cell unfolded protein response and apoptosis. Hum Mol Genet. 2012;21(2):384–393. doi:10.1093/hmg/ddr473

37. Ogando DG, Shyam R, Kim ET, Wang YC, Liu CY, Bonanno JA. Inducible Slc4a11 Knockout Triggers Corneal Edema Through Perturbation of Corneal Endothelial Pump. Invest Ophthalmol Vis Sci. 2021;62(7):28. doi:10.1167/iovs.62.7.28

38. Shyam R, Ogando DG, Bonanno JA. Mitochondrial ROS in Slc4a11 KO Corneal Endothelial Cells Lead to ER Stress. Front Cell Dev Biol. 2022;10:878395. doi:10.3389/fcell.2022.878395

39. Gendron SP, Thériault M, Proulx S, Brunette I, Rochette PJ. Restoration of Mitochondrial Integrity, Telomere Length, and Sensitivity to Oxidation by In Vitro Culture of Fuchs’ Endothelial Corneal Dystrophy Cells. Investigative Ophthalmology & Visual Science. 2016;57(14):5926–5934. doi:10.1167/iovs.16-20551

40. Czarny P, Seda A, Wielgorski M, et al. Mutagenesis of mitochondrial DNA in Fuchs endothelial corneal dystrophy. Mutation Research/Fundamental and Molecular Mechanisms of Mutagenesis. 2014;760:42–47. doi:10.1016/j.mrfmmm.2013.12.001

41. Halilovic A, Schmedt T, Benischke AS, et al. Menadione-Induced DNA Damage Leads to Mitochondrial Dysfunction and Fragmentation During Rosette Formation in Fuchs Endothelial Corneal Dystrophy. Antioxidants & Redox Signaling. 2016;24(18):1072–1083. doi:10.1089/ars.2015.6532

42. Jurkunas UV, Bitar MS, Funaki T, Azizi B. Evidence of oxidative stress in the pathogenesis of fuchs endothelial corneal dystrophy. Am J Pathol. 2010;177(5):2278–2289. doi:10.2353/ajpath.2010.100279

43. Katikireddy KR, White TL, Miyajima T, et al. NQO1 downregulation potentiates menadione-induced endothelial-mesenchymal transition during rosette formation in Fuchs endothelial corneal dystrophy. Free Radic Biol Med. 2018;116:19–30. doi:10.1016/j.freeradbiomed.2017.12.036

44. Guha S, Chaurasia S, Ramachandran C, Roy S. SLC4A11 depletion impairs NRF2 mediated antioxidant signaling and increases reactive oxygen species in human corneal endothelial cells during oxidative stress. Sci Rep. 2017;7(1):4074. doi:10.1038/s41598-017-03654-4

45. Han SB, Ang HP, Poh R, et al. Mice With a Targeted Disruption of Slc4a11 Model the Progressive Corneal Changes of Congenital Hereditary Endothelial Dystrophy. Investigative Ophthalmology & Visual Science. 2013;54(9):6179–6189. doi:10.1167/iovs.13-12089

46. Ogando DG, Choi M, Shyam R, Li S, Bonanno JA. Ammonia sensitive SLC4A11 mitochondrial uncoupling reduces glutamine induced oxidative stress. Redox Biol. 2019;26:101260. doi:10.1016/j.redox.2019.101260

47. Kumar V, Jurkunas UV. Mitochondrial Dysfunction and Mitophagy in Fuchs Endothelial Corneal Dystrophy. Cells. 2021;10(8):1888. doi:10.3390/cells10081888

48. Hogan M, Alvarado J, Weddell J. Histology ofthe human eye. Philadelphia, W B Saunders. Published online 1971:269.

49. Damkier HH, Nielsen S, Praetorius J. Molecular expression of SLC4-derived Na+dependent anion transporters in selected human tissues. Am J Physiol Regul Integr Comp Physiol. 2007;293(5):R2136–2146. doi:10.1152/ajpregu.00356.2007

50. Choi M, Bonanno JA. Mitochondrial Targeting of the Ammonia-Sensitive Uncoupler SLC4A11 by the Chaperone-Mediated Carrier Pathway in Corneal Endothelium. Invest Ophthalmol Vis Sci. 2021;62(12):4. doi:10.1167/iovs.62.12.4

51. Li S, Kim E, Ogando DG, Bonanno JA. Corneal Endothelial Pump Coupling to Lactic Acid Efflux in the Rabbit and Mouse. Invest Ophthalmol Vis Sci. 2020;61(2):7. doi:10.1167/iovs.61.2.7

52. Elhalis H, Azizi B, Jurkunas UV. Fuchs endothelial corneal dystrophy. Ocul Surf. 2010;8(4):173–184. doi:10.1016/s1542-0124(12)70232-x

53. Laing RA, Sanstrom MM, Berrospi AR, Leibowitz HM. Changes in the corneal endothelium as a function of age. Exp Eye Res. 1976;22(6):587–594. doi:10.1016/0014-4835(76)90003-8

54. Cheng H, Jacobs PM, McPherson K, Noble MJ. Precision of cell density estimates and endothelial cell loss with age. Arch Ophthalmol. 1985;103(10):1478–1481. doi:10.1001/archopht.1985.01050100054017

55. Abib FC, Barreto Junior J. Behavior of corneal endothelial density over a lifetime. J Cataract Refract Surg. 2001;27(10):1574–1578. doi:10.1016/s0886-3350(01)00925-7

56. Joyce NC. Proliferative capacity of corneal endothelial cells. Exp Eye Res. 2012;95(1):16–23. doi:10.1016/j.exer.2011.08.014

57. McCarey BE, Edelhauser HF, Lynn MJ. Review of corneal endothelial specular microscopy for FDA clinical trials of refractive procedures, surgical devices, and new intraocular drugs and solutions. Cornea. 2008;27(1):1–16. doi:10.1097/ICO.0b013e31815892da

58. Mohammad-Salih P a. K. Corneal endothelial cell density and morphology in normal Malay eyes. Med J Malaysia. 2011;66(4):300–303.

59. Vassilev VS, Mandai M, Yonemura S, Takeichi M. Loss of N-Cadherin from the Endothelium Causes Stromal Edema and Epithelial Dysgenesis in the Mouse Cornea. Investigative Ophthalmology & Visual Science. 2012;53(11):7183–7193. doi:10.1167/iovs.12-9949

60. Thériault M, Gendron SP, Brunette I, Rochette PJ, Proulx S. Function-Related Protein Expression in Fuchs Endothelial Corneal Dystrophy Cells and Tissue Models. The American Journal of Pathology. 2018;188(7):1703–1712. doi:10.1016/j.ajpath.2018.03.014

61. Srinivas SP. Cell Signaling in Regulation of the Barrier Integrity of the Corneal Endothelium. Exp Eye Res. 2012;95(1):8–15. doi:10.1016/j.exer.2011.09.009

62. Herce-Pagliai C, Kotecha S, Shuker DEG. Analytical Methods for 3-Nitrotyrosine as a Marker of Exposure to Reactive Nitrogen Species: A Review. Nitric Oxide. 1998;2(5):324–336. doi:10.1006/niox.1998.0192

63. Bandookwala M, Sengupta P. 3-Nitrotyrosine: a versatile oxidative stress biomarker for major neurodegenerative diseases. Int J Neurosci. 2020;130(10):1047–1062. doi:10.1080/00207454.2020.1713776

64. Shyam R, Ogando DG, Choi M, Liton PB, Bonanno JA. Mitochondrial ROS Induced Lysosomal Dysfunction and Autophagy Impairment in an Animal Model of Congenital Hereditary Endothelial Dystrophy. Invest Ophthalmol Vis Sci. 2021;62(12):15. doi:10.1167/iovs.62.12.15

65. Piccolo S, Dupont S, Cordenonsi M. The Biology of YAP/TAZ: Hippo Signaling and Beyond. Physiological Reviews. 2014;94(4):1287–1312. doi:10.1152/physrev.00005.2014

66. Leonard BC, Park S, Kim S, et al. Mice Deficient in TAZ (Wwtr1) Demonstrate Clinical Features of Late-Onset Fuchs’ Endothelial Corneal Dystrophy. Invest Ophthalmol Vis Sci. 2023;64(4):22. doi:10.1167/iovs.64.4.22

67. Liu C, Miyajima T, Melangath G, et al. Ultraviolet A light induces DNA damage and estrogen-DNA adducts in Fuchs endothelial corneal dystrophy causing females to be more affected. Proceedings of the National Academy of Sciences. 2020;117(1):573–583. doi:10.1073/pnas.1912546116

68. Han S, Mueller C, Wuebbolt C, et al. Selective effects of estradiol on human corneal endothelial cells. Sci Rep. 2023;13(1):15279. doi:10.1038/s41598-023-42290-z

69. Price MO, Mehta JS, Jurkunas UV, Price FW. Corneal endothelial dysfunction: Evolving understanding and treatment options. Progress in Retinal and Eye Research. 2021;82:100904. doi:10.1016/j.preteyeres.2020.100904

70. Yong KL, Nguyen HV, Cajucom-Uy HY, et al. Cost Minimization Analysis of Precut Cornea Grafts in Descemet Stripping Automated Endothelial Keratoplasty. Medicine (Baltimore). 2016;95(8):e2887. doi:10.1097/MD.0000000000002887

71. Kinoshita S, Koizumi N, Ueno M, et al. Injection of Cultured Cells with a ROCK Inhibitor for Bullous Keratopathy. New England Journal of Medicine. 2018;378(11):995–1003. doi:10.1056/NEJMoa1712770

72. Numa K, Imai K, Ueno M, et al. Five-Year Follow-up of First 11 Patients Undergoing Injection of Cultured Corneal Endothelial Cells for Corneal Endothelial Failure. Ophthalmology. 2021;128(4):504–514. doi:10.1016/j.ophtha.2020.09.002

73. Aiello F, Gallo Afflitto G, Ceccarelli F, Cesareo M, Nucci C. Global Prevalence of Fuchs Endothelial Corneal Dystrophy (FECD) in Adult Population: A Systematic Review and Meta-Analysis. J Ophthalmol. 2022;2022:3091695. doi:10.1155/2022/3091695

